# A tissue-like platform for studying engineered quiescent human T-cells’ interactions with dendritic cells

**DOI:** 10.1101/587386

**Authors:** Enas Abu-Shah, Philippos Demetriou, Stefan Balint, Viveka Mayya, Mikhail A. Kutuzov, Omer Dushek, Michael L. Dustin

## Abstract

Research in the field of human immunology is restricted by the lack of a system that reconstitutes the *in-situ* activation dynamics of quiescent human antigen-specific T-cells interacting with dendritic cells. Here we report a tissue-like system that recapitulates the dynamics of engineered primary human immune cell. Our approach facilitates real-time single cell manipulations, tracking of interactions and functional responses complemented by population-based measurements of cytokines, activation status and proliferation. As a proof of concept, we recapitulate immunological phenomenon such as CD4 help to CD8 T-cells through enhanced maturation of DCs and effect of PD-1 checkpoint blockades. In addition, we characterise unique dynamics of T-cell/DC interactions as a function of antigen affinity.

## Introduction

The field of human immunology has gained much attention in the last few decades with the increasing realisation that despite the tremendous utility of mouse models, they can only provide limited insight for advancing translational research ^1, 2^. Most of the research in human immunology is based on observational studies, whereby, for example, data sets of gene expression profiles are used to identify disease biomarkers. Although these studies promote our understanding and help in developing hypotheses about disease mechanisms, they fall short, and experimental research is needed ^3^. The classical tools in experimental human immunology rely heavily on immortalised cell lines, stimulated clonal lines, peripheral blood, and more rarely humanised mouse models. Each of these tools has its limitation. Cell lines have diverged tremendously from primary cells, making their signalling machinery and response thresholds no longer representative ^4^; bulk peripheral blood assays lack the spatial organisation of tissues; and humanised mouse models have only a portion of their system “humanised”, rendering the remaining parts confounding factors ^5^. To address these limitations, we introduce a holistic approach to allow researchers to manipulate and study human immune cells in a tissue-like environment with minimal aberrations to their natural behaviours. We report a toolkit that achieves two critical goals for engineering T-cells: (1) high efficiency TCR expression and maintenance of T-cell motility, activation dynamics and viability using mRNA electroporation and (2) a tissue-like 3D culture (Figure 1).

**Figure 1.**
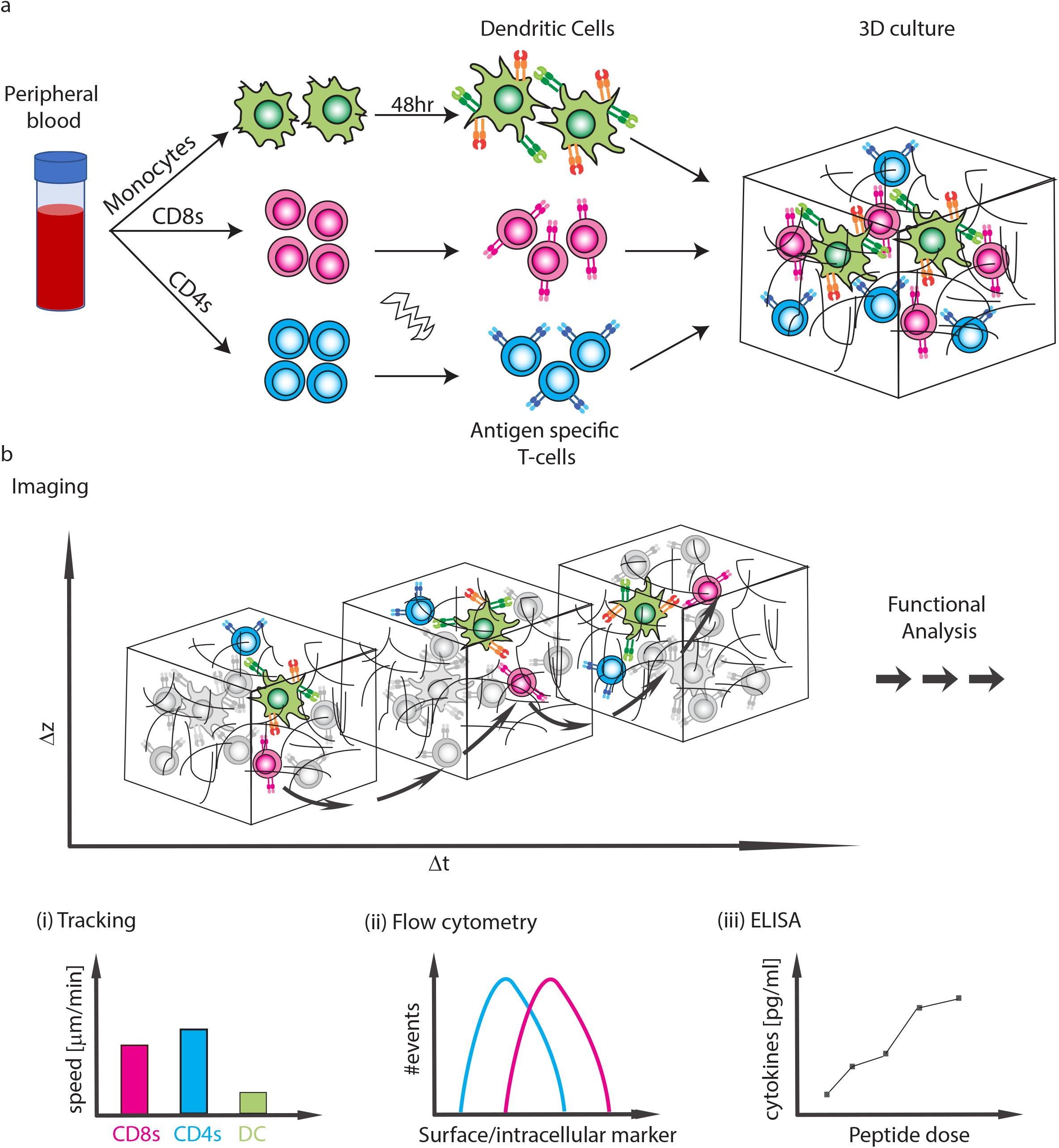
Schematics of experimental system. (a) Isolation of CD4 T-cells, CD8 T-cells and monocytes from peripheral blood. Subpopulations of T-cells can be further isolated and TCR expression is induced by mRNA electroporation. Monocytes are induced to differentiate into DCs, which are also activated with a 48 hr express protocol. The cells are then moved into collagen gel-based 3D culture. (b) Cells can be taken for imaging or downstream functional analyses. (i) The imaging can be used to analyse interactions dynamics such as speed or conjugate formation, (ii) Cells can be extracted for flow cytometry, (iii) Cytokine production into the culture can be measured.

## Results

### Engineering quiescent primary T-cells

Successful engineering of T-cell specificity requires the expression of a functional and stable exogenous TCR, as well as the preservation of important cellular characteristics. One such characteristic is cell motility, which is essential for efficient antigen search in a physiological 3D setting. Here, we have optimised the procedures to engineer naïve human CD4 and CD8 T-cells using mRNA electroporation. The common approach to render T-cells antigen specific relies on lentiviral transduction of the TCR which requires prior expansion of T-cells to achieve high expression efficiency and hinders the study of natural quiescent populations, namely naïve and memory T-cells ^6^. To avoid the need for cell expansion, it is possible to use electroporation to deliver the genetic materials. However, reported approaches using electroporation suffer from low efficiency of expression, low viability, and most importantly impaired cell motility, a critical consideration in the context of delivery to target tissues. Using a square-wave electroporation apparatus, we are able to express the **1G4** TCR, specific to the NY-ESO tumour associated antigen^7^ in naïve (Figure 2A) and memory (Figure 2- figure supplement 1A) CD8 T-cells as well as the **868** TCR specific for the HIV gag protein ^8^ (Figure 2- figure supplement 1B), both presented by HLA-A*0201. Successful pairing of the alpha and beta heterodimer of the TCR is important for the expression and assembly at the cell surface, this was achieved by introducing cysteine modifications in the TCR chains (Figure 2- figure supplement 1B and Table 1) ^9^, as well as co-transfection with the human CD3ζ (Figure 2- figure supplement 1C), as the endogenous protein is most likely sequestered by the endogenous TCR limiting the expression of the introduced TCR. The functionality of naïve CD8 T-cells following TCR electroporation was confirmed by the formation of a mature immunological synapse in naïve and activated CD8 T-cells (IS, Figure 2B and Figure 2- figure supplement 1D-E), and the upregulation of the activation marker CD69 (Figure 2- figure supplement 2A). The efficiency of TCR expression varied among donors with a mean value of 81 ± 7% (mean±SD, n=13), with more than 90% cell viability and 80-90% cell recovery, both of which were severely reduced using alternative electroporation approaches (Figure 2- figure supplement 3).

**Figure 2.**
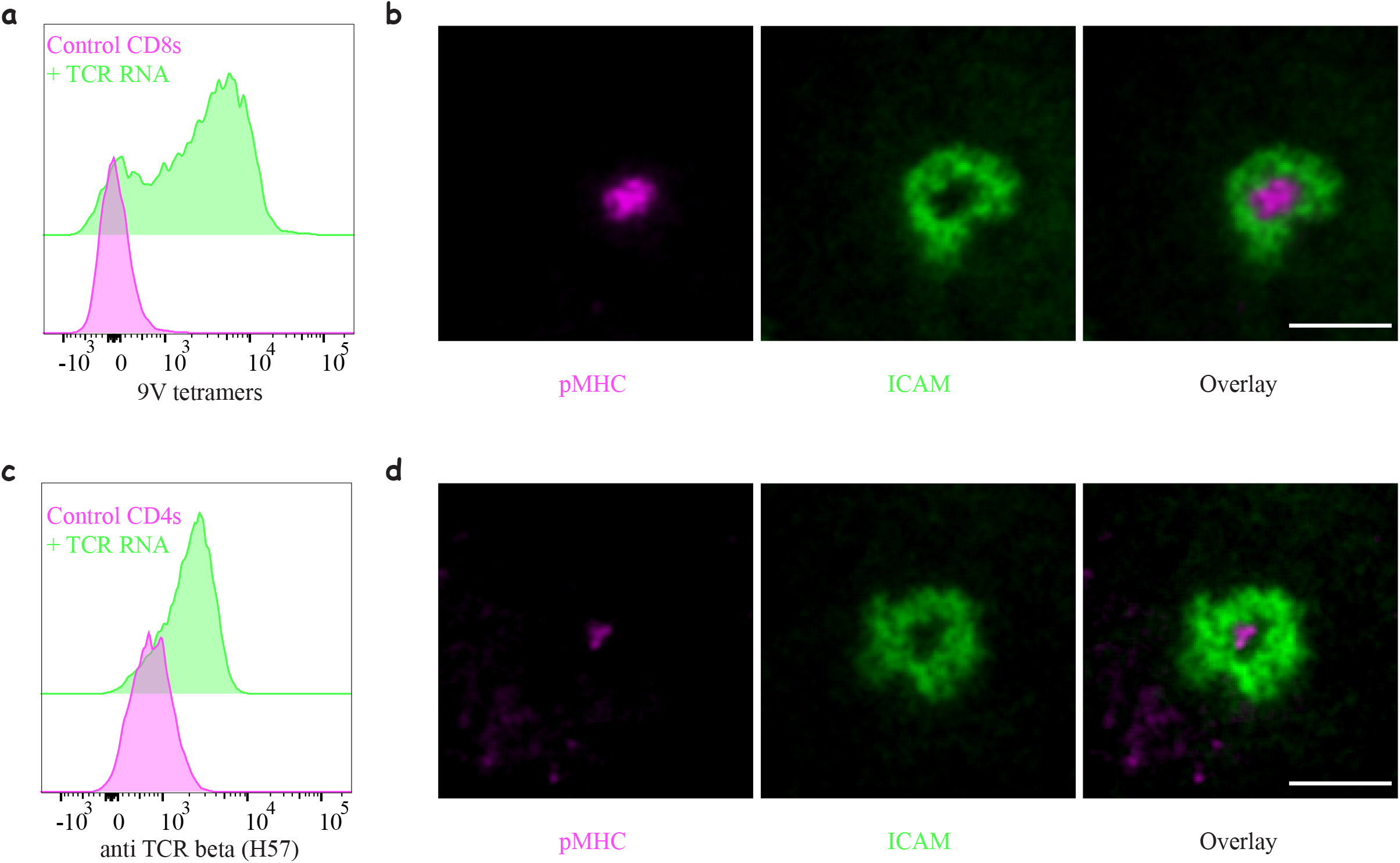
Engineering Immune Cells for Antigen-specific Interactions. (a) Expression of **1G4** TCR in naïve CD8 T-cells upon mRNA electroporation detected using NY-ESO-9V/HLA-A2 tetramer, ~80% positive (Representative of N=13). (b) Formation of an immunological synapse by **1G4**-expressing naïve CD8 T-cells on supported lipid bilayers (SLBs) with cSMAC enriched with 9V/A2 pMHC (magenta) surrounded by LFA/ICAM1 ring (green). Representative of at least 3 independent repeats. (c) Expression of **6F9** TCR in naïve CD4 T-cells detected using an antibody against the constant region of mouse TCRβ, ~67% positive (Representative of N=15). (d) Formation of immunological synapse by **6F9**-expressing naïve CD4 T-cells on SLB containing MAGE/DP4 pMHC (magenta). Representative of at least 3 independent repeats. Scale bars = 5 μm.

**Table 1:** Sequence modification of MHC-I restricted TCRs.

| TCR chain | Original sequence | Modified sequence |
| --- | --- | --- |
| Alpha | DKTVL | DKCVL |
| Beta | GVSTD | GVCTD |

The induction of expression of an exogenous TCR in CD4 T-cells is considerably harder than in CD8 T-cells ^6^. Using natural sequences or the introduction of cysteine modifications used for **1G4** and **868** failed to induce expression of TCRs in naïve CD4 T-cells. However, replacing the constant region of the human TCR with that of mouse TCR (Table 2) ^10^ allowed for successful chain pairing and we were able to achieve expression in 72 ± 8.9% (mean±SD, n=15) of the cells detected using H57 mAb against the mouse Cβ. We have successful expressed three different HLA-DPB1*04 restricted TCRs in naïve CD4s: **6F9** (Figure 2C) and **R12C9**, both specific to the MAGE-A3 protein ^11^ and **SG6** specific for NY-ESO ^12^ (Figure 2- figure supplement 4A). The expression of **SG6** was consistently lower than **6F9** and **R12C9** in naïve CD4 T-cells. However, when the same TCRs were electroporated into previously activated CD4 T-cells, high expression levels were achieved in 97.8 ± 1% (mean±SD, n=15) of the cells (Figure 2- figure supplement 4B). The functionality of the naïve CD4 T-cells upon TCR electroporation was confirmed by IS formation (Figure 2D and Figure 2- figure supplement 4C-D) and T-cell activation (Figure 2- figure supplement 2B). The enhanced expression of the TCR in activated CD4 T-cells resulted in enhanced pMHC accumulation at the IS (Figure 2- figure supplement 4D).

**Table 2:**
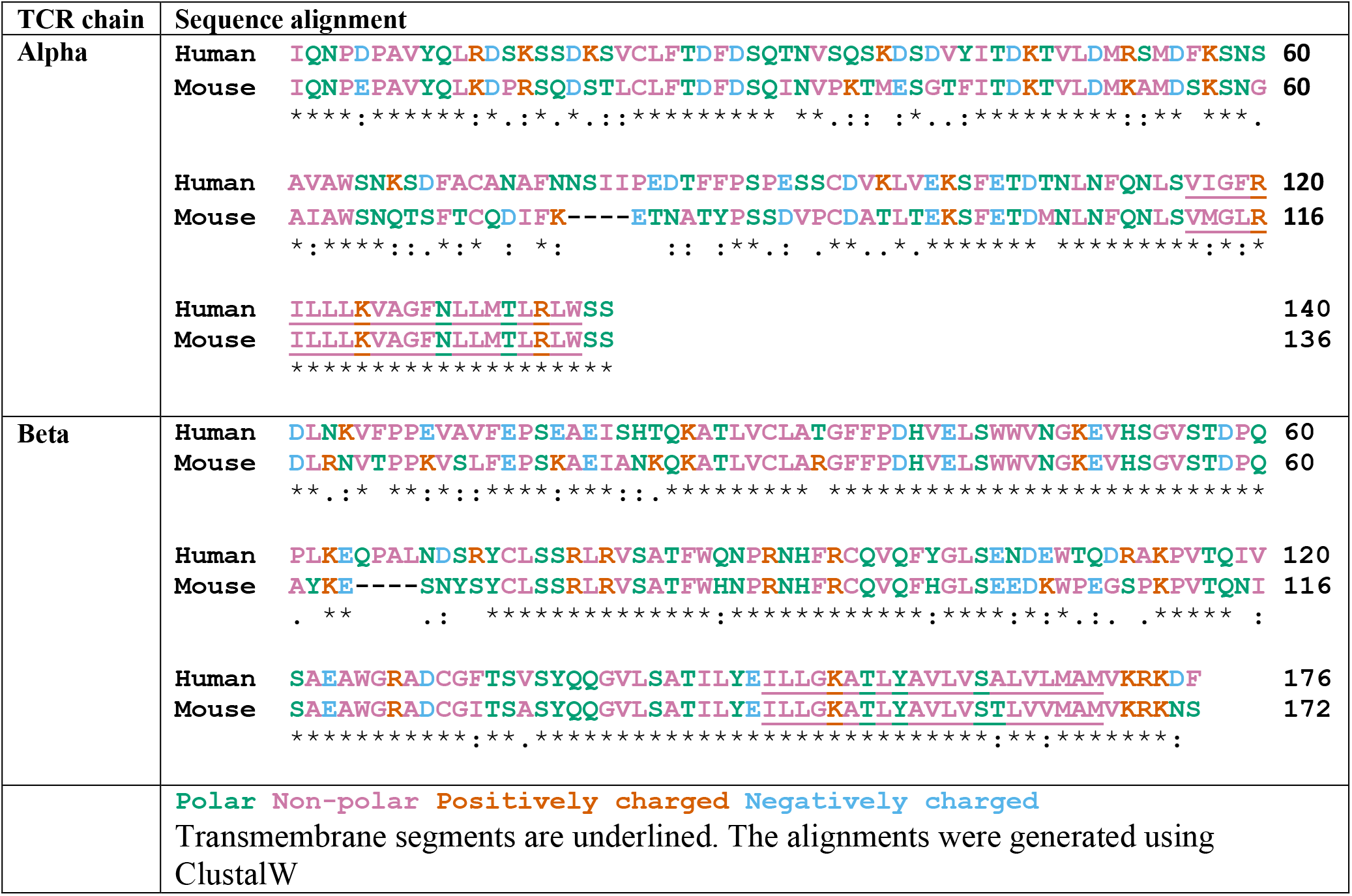
Sequence alignment of human and mouse constant region used to modify MHC-II restricted TCRs.

Using the square-wave electroporator we were able to maintain the motile behaviour and interaction dynamics of naïve CD8 T-cells expressing **1G4**, at similar levels to those of untouched cells, on 2D stimulatory ‘spots’ (Figure 2- figure supplement 5, Video 1 and ^13^).

### Express monocyte-derived Dendritic cells as model Antigen presenting cells

Naïve T-cells get activated by interacting with professional antigen presenting cells (APCs); the most potent are dendritic cells (DCs). The closest model for *bona fide* DCs is monocyte-derived DCs (moDCs) with well-established protocols for their generation. The caveat with those protocols is their extended duration (7-10 days) ^14^, a time-frame which may alter autologous T-cells’ phenotype. We therefore optimised a 48 h protocol ^15^ to generate fully matured moDCs. During the first 24 h monocyte differentiation into DC was induced, followed by 24 h maturation to get activated DCs (acDC) (Figure 3). The “express” moDCs are able to upregulate MHC and other costimulatory molecules and produce a similar profile of soluble factors as the “classical” moDC (Figure 3- figure supplement 1A-B). We have also quantified peptide loading on these cells and achieved levels similar to those previously reported with other APCs (Figure 3- figure supplement 1C) ^16^.

**Figure 3.**
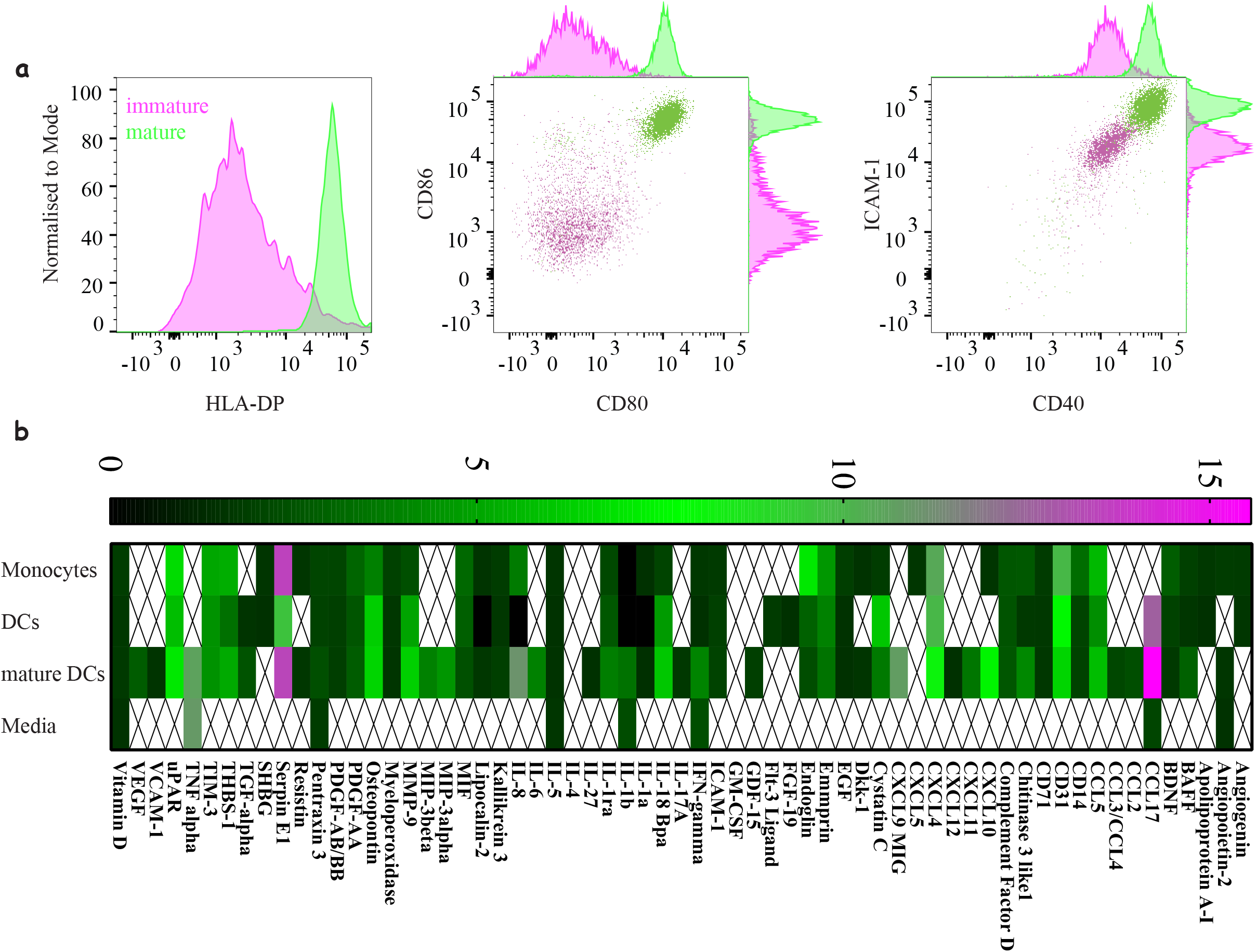
Characterising “Express” Dendritic as a model antigen presenting cell. (a) Activation and differentiation profile of “express” monocyte-derived dendritic cells: mature cells (green) upregulate their antigen presentation and costimulatory molecules compared to immature cells (magenta). Representative of at least 3 independent repeats. (b) Cytokine and chemokine secretion profile from monocyte, dendritic cells and mature dendritic cells using the 48hr express protocol. An average of three donors where signals bellow 1.5-fold above background were not included.

### Collagen-based 3D model to support immune cell migration and interactions

T-cells and DCs interactions have been extensively studied in mice using explants and intravital imaging ^17^, such studies are practically impossible in humans. Furthermore, accurate control and manipulation of parameters such as antigen dose and cell ratios is limited. Having successfully engineered naïve human T-cells we set out to establish a flexible 3D platform to interrogate their dynamics and interactions with APC using a multitude of correlated functional readouts. The desired platform should: (1) support the culture of T-cells and APCs for multiple days, (2) support the motility of the cells, (3) allow live imaging, (4) enable downstream analysis and (5) be easy to use. To achieve these goals we chose collagen I based 3D matrices^18^, which we optimised to support the culture of human immune cells. Culturing cells in the presence of human serum (HS) results in better basal motility than FBS (Figure 4A, Figure 4- figure supplement 1A and Video 2, Table 3), which is significantly enhanced by the addition of homeostatic chemokines such as CCL19 (Video 3), CCL21 and CXCL12 (Video 4). However, only CXCL12 is able to maintain high motility in cultures with FBS. We have explored different sources of collagen I (bovine, human and non-tryptic collagen) and other complex extracellular matrices, ECM (Figure 4- figure supplement 1B). All collagens were equally good in supporting motility and interactions, while more advanced ECM may have a marginal advantage for motility. Yet, the larger batch variation, higher background activation and the complexity in extracting cells for downstream analysis weighed against adopting them for our assays. In our view, the use of bovine collagen-I as the 3D scaffold, with human serum, and CXCL12 to enhance motility provides an optimal 3D system with similar movement parameters to human T cells in mouse lymph nodes or explanted human tissues ^19, 20, 21, 22^. We have also re-evaluated two additional electroporation methods, namely Lonza-Amaxa and ThermoFisher-Neon, In addition to their lower performance in cell recovery and protein expression (Figure 2- figure supplement 3) they further resulted in poor motility dynamics and hence activation of cells within the 3D environment, where the search for antigen loaded APCs becomes a confounding factor (Figure 4- figure supplement 1C-F).

**Figure 4.**
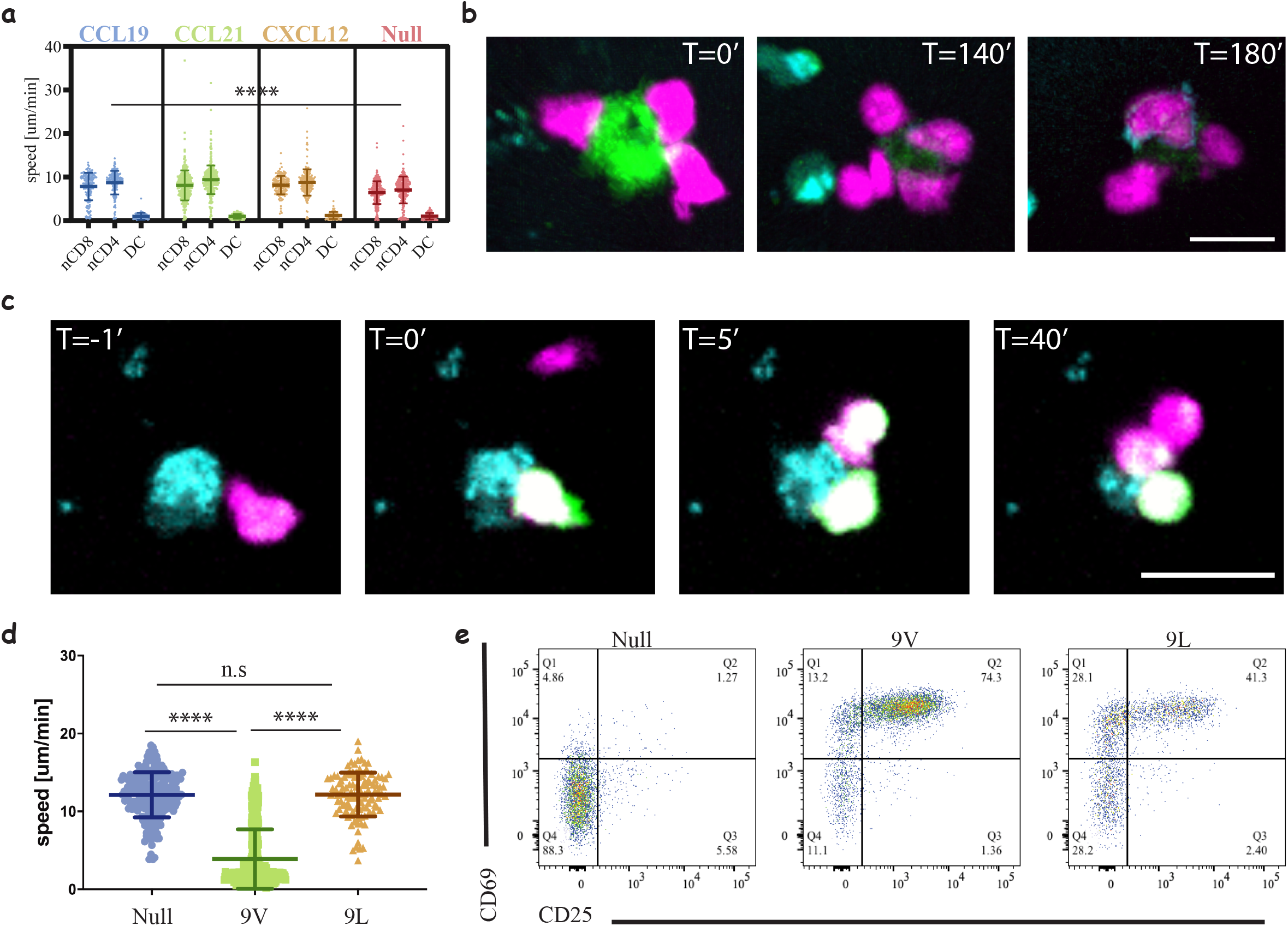
Three-Dimensional Culture System to Study Immune Cell Interactions. (a) Motility speeds of naïve CD4 and CD8 T-cells and acDCs in collagen gels with different chemokines. Note the faster motility of T-cells compared to acDC and equally efficient motility using homeostatic chemokines (CCL19, CCL21 and CXCL12). Representative of 2 independent repeats (Video 2, Table 3). (b) Snapshots from a time-lapse movie (Video 3) following the interactions of **1G4**-expressing naïve CD8 T-cells (magenta) with antigen loaded acDC (green) in collagen gels containing CXCL12, and the upregulation of CD69 is visible at T=140 min following the accumulation of an anti-CD69 antibody on the surface of the cells (cyan, also in cyan are irrelevant CD4 T-cells). Representative of 3 independent repeats. (c) Snapshots from a time-lapse movie (Video 4) showing three different **1G4**-expressing CD8 T-cells (magenta), loaded with calcium dye Fluo4-AM (green), interacting with antigen loaded acDCs (100 nM NY-ESO-9V, cyan) in 3D collagen and fluxing calcium (green) upon binding. Note the lower flux in the second and third contacts suggesting lower antigen presentation sequestered by the primary synapse. (d) Speed of **1G4**-expressing naïve CD8 T-cells upon culture in collagen gels containing CXCL12, with acDC either loaded with high affinity (100 nM NY-ESO-9V) or low affinity (100 nM NY-ESO-9L) peptide or unloaded (null). Note the deceleration of the cells upon engaging with high affinity peptide (3 μm/min compared to 12 μm/min for both null and 9L). Representative of 2 independent repeats. (Videos 7-9). (d) The cells from (c) extracted from the collagen gel and run on a flow cytometry to look at CD69 and CD25 as activation markers. Note the good activation despite the absence of T-cell arrest with low affinity peptide (NY-ESO-9L). Representative of more than 3 independent repeats. Scale bars = 10 μm. (****, p < 0.0001, ANOVA)

**Table 3:**
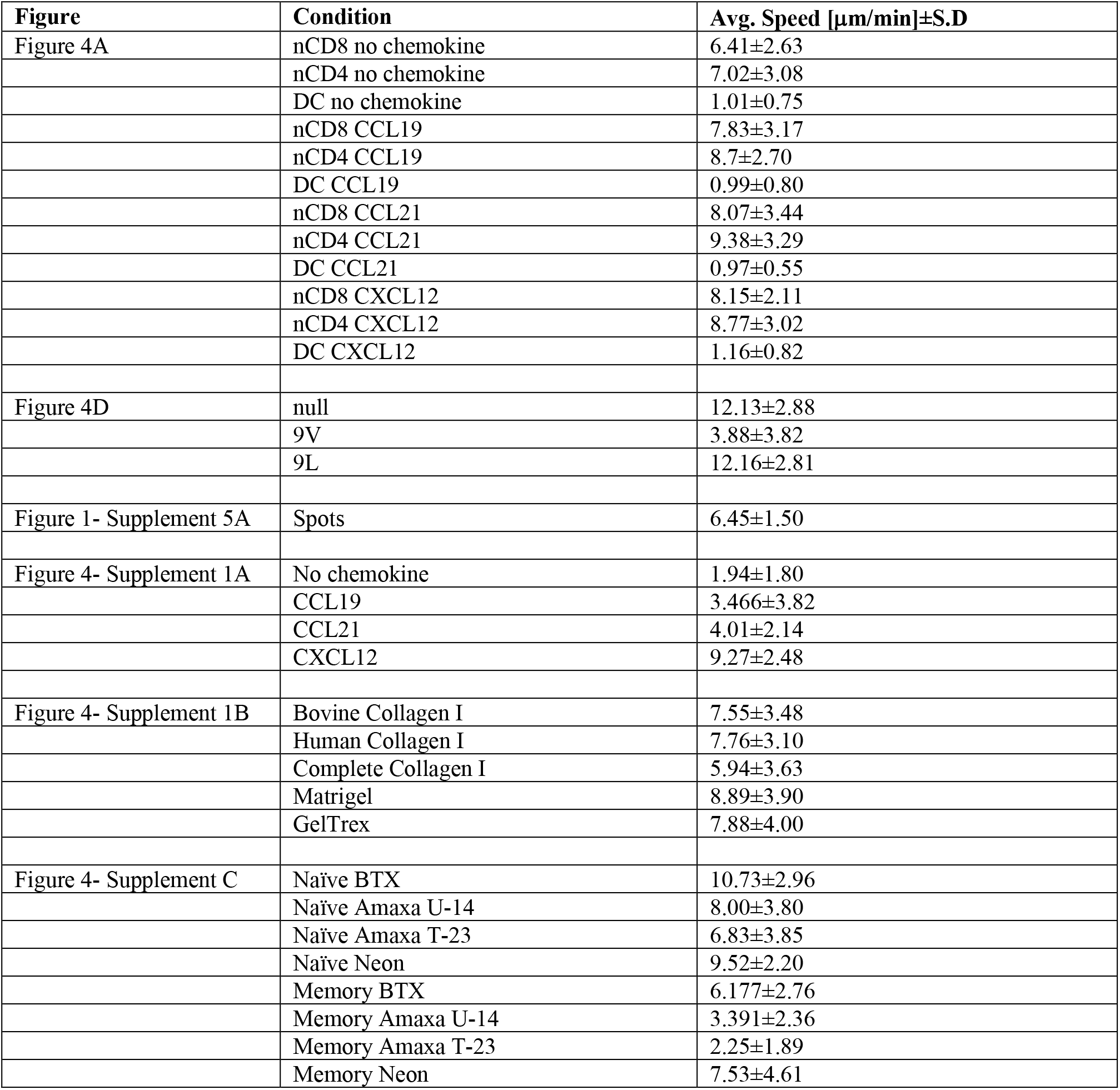
Average speeds for the plots in the figures.

| Figure | Condition | Avg. Speed [$\mu\text{m}/\text{min}$]±S.D |
| --- | --- | --- |
| Figure 4A | nCD8 no chemokine | 6.41±2.63 |
| | nCD4 no chemokine | 7.02±3.08 |
| | DC no chemokine | 1.01±0.75 |
| | nCD8 CCL19 | 7.83±3.17 |
| | nCD4 CCL19 | 8.7±2.70 |
| | DC CCL19 | 0.99±0.80 |
| | nCD8 CCL21 | 8.07±3.44 |
| | nCD4 CCL21 | 9.38±3.29 |
| | DC CCL21 | 0.97±0.55 |
| | nCD8 CXCL12 | 8.15±2.11 |
| | nCD4 CXCL12 | 8.77±3.02 |
| | DC CXCL12 | 1.16±0.82 |
| Figure 4D | null | 12.13±2.88 |
| | 9V | 3.88±3.82 |
| | 9L | 12.16±2.81 |
| Figure 1- Supplement 5A | Spots | 6.45±1.50 |
| Figure 4- Supplement 1A | No chemokine | 1.94±1.80 |
| | CCL19 | 3.466±3.82 |
| | CCL21 | 4.01±2.14 |
| | CXCL12 | 9.27±2.48 |
| Figure 4- Supplement 1B | Bovine Collagen I | 7.55±3.48 |
| | Human Collagen I | 7.76±3.10 |
| | Complete Collagen I | 5.94±3.63 |
| | Matrigel | 8.89±3.90 |
| | GelTrex | 7.88±4.00 |
| Figure 4- Supplement C | Naïve BTX | 10.73±2.96 |
| | Naïve Amaxa U-14 | 8.00±3.80 |
| | Naïve Amaxa T-23 | 6.83±3.85 |
| | Naïve Neon | 9.52±2.20 |
| | Memory BTX | 6.177±2.76 |
| | Memory Amaxa U-14 | 3.391±2.36 |
| | Memory Amaxa T-23 | 2.25±1.89 |
| | Memory Neon | 7.53±4.61 |

### Low affinity peptides result in T-cell activation without stable immunological synapse

We present here a proof of concept to the usefulness of our set-up by interrogating naïve CD8 T-cells activation. We used live cell imaging to follow the interactions in real-time by upregulation of CD69 (Figure 4B and Video 5) as well as calcium flux (Figure 4C and Video 6). We observed major changes in interaction dynamics as a function of peptide affinity ^23^ (Figure 4D). A high affinity peptide, NY-ESO-9V, (Video 7) induced T-cell arrest, whereas a lower affinity peptide, NY-ESO-9L, (Video 8), led to similar dynamics as a null peptide (Video 9) where no stable contacts are formed. Since our 3D culture offers a relatively simple way to extract cells for flow cytometry and downstream analyses, we were able to correlate the interaction dynamics with the activation state of the cells and observe marginal differences in the response relative to the major differences in dynamics (Figure 4E, Figure 4- supplement figure 1A). Using our system, we are also able to monitor cytokine production (Figure 4- supplement figure 1B), proliferation (Figure 4- supplement figure 1C) and intracellular activation markers (Figure 4- supplement figure 1D). In order to study the interaction dynamics of T-cells at different time points we can pre-load the DCs prior or following gel polymerisation to mimic the delivery of antigen to lymph nodes (Figure 4- supplement figure 1E and Videos 10-11, respectively).

### Enhancing CD8 responses through CD4 help and PD1 checkpoint blockade

Our ability to engineer both CD4 and CD8 T-cells independently with different TCRs enables us to interrogate complex dynamics of trinary cellular system whereby both types of T-cells can interact with similar or different APCs. We show here that using our system we can interrogate the dynamics of CD4 T-cells help to CD8 T-cells ^24^ (Figure 5A, Video 12) where the activation of CD4 T-cells (Figure 5B, top) coincides with enhanced maturation of the DCs by upregulation of CD86 (Figure 5B, middle) and enhanced activation of the CD8 T-cells (Figure 5B, bottom).

**Figure 5.**
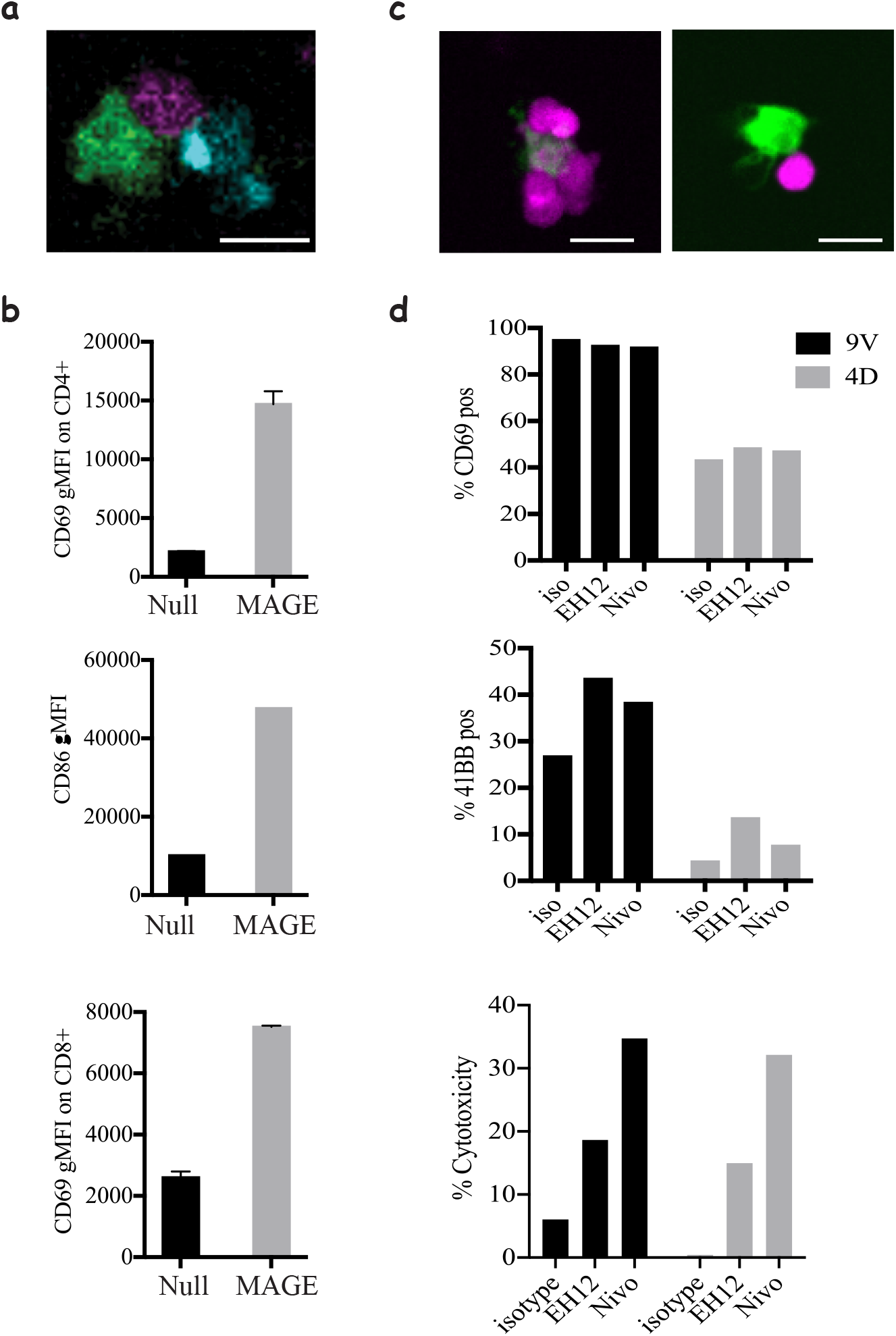
Modelling CD4 help and immune checkpoint in 3D cultures. (a) Co-culture of **1G4**-expressing naïve CD8 T-cells (magenta), **6F9**-expressing naïve CD4 T-cells (cyan) and acDC loaded with 100nM NY-ESO-6T and 10μM MAGE-A3_243-258_ peptides (green), (Video 12), in collagen gels containing CXCL12. (b) Upregulation of CD69 on CD4 (top), and CD8 T-cells (bottom) and the upregulation of the costimulatory molecule CD86 on acDCs (middle) in the co-culture system from (a) in the presence and absence of MAGE as a stimulus for CD4 T-cells. Representative of at least 3 independent repeats. In all experiments the cell ratio was (5:5:1, CD4: CD8: acDC). (c) **1G4**-expressing memory CD8 T-cells (magenta) co-cultured with acDC loaded with 100nM NY-ESO-9V (green). Cells form cluster around acDC in the presence of anti PD-1 blocking antibody Nivolumab (left) compared to less tight contacts in the isotype control (right). (d) The activation of the CD8 T-cells by upregulation of CD69 (top) and 41BB (middle) as well as cytotoxicity measured as the release of LDH following the killing of the target (bottom). In all experiments the cell ratio was (10:1, CD8: acDC). Scale bars = 10 μm.

Furthermore, we utilised our system to mimic one of the most successful immune checkpoint blockades currently in clinical use; PD-1 blockade, which is known to act by enhancing CD8 T-cell responses towards target cells. To that end, we used memory CD8 T-cells and treated them with two clones of blocking antibodies against PD1; Nivolumab and EH12. Despite the lack of any change in the motility dynamics of the cells (Figure 5- Supplement Figure 1A), there was a clear evidence that the presence of the checkpoint blockade enhances the clustering of CD8 T-cells around their targets (Figure 5C). This has also been corroborated by enhanced activation, although only observable for late readouts such as 41BB expression and target cell killing (Figure 5D). Interestingly we saw an enhanced expression of PDL-1 (Figure 5- Supplement Figure 1B), the ligand for PD-1, on CD8 T-cells, which could lead to cis interactions that act as a cell intrinsic regulation mechanism ^25^. Surprisingly, we noted that there are differences between the pathways enhanced by the two blocking antibody clones where EH12 seems superior at enhancing surface markers, whereas Nivolumab outperforms EH12 in the killing assays.

## Discussion

In this report, we present, to the best of our knowledge, a unique and first of its kind holistic approach to engineer quiescent antigen specific human T-cells and study them in a context akin to physiological settings. Our tools are widely applicable to generate T-cells expressing different receptors and other proteins with high efficiency and minimal adverse effects associated with other engineering methods, which are essential for implementation of a successful translational approach. We provide a proof of concept where our system is used to probe the dynamics of T-cell/APC interactions as a function of peptide affinity, recapitulate CD4 help to CD8 T-cells and interrogate PD1 checkpoint blockade. Our experimental approach can be further extended towards reconstitution of more complex biology such as tumour infiltrating lymphocytes and the role of regulatory T-cells in cancer. We believe that the mechanical and biophysical features introduced by our platform provide greater fidelity to *in-vivo* conditions and will improve predictive power of pre-clinical studies in all human cell-based systems. We propose that our system should be integrated with other classical and computational approaches to enhance translational research.

## Acknowledgements

We thank Vincenzo Cerundulo for the **1G4** TCR sequence, Andrew Sewell for the **868** TCR plasmid, Steven Rosenberg for the **6F9**, **R12C9** and **SG6** TCR sequences and James Riley for pGEM-64A plasmid and introducing us to mRNA electroporation. We thank Jehan Afrose and Heather Rada for the generation of recombinant proteins. We thank the NIH tetramer facility for providing pMHC monomers. We thank Arbel Artzy-Schnirman for critical reading of the manuscript and help with graphical figures. We thank Kinneret Keren, Erez Braun, Anton van der Merwe, Marion H. Brown and David Depoil for discussions and feedback.

The work has been supported by an UCB-Oxford Post-doctoral Fellowship to E.A-S., European Research Council ERC-2014-AdG_670930 for V.M. and S.B., Kennedy Trust for Rheumatology (KTRR) Prize Studentship to P.D., a Wellcome Trust Principal Research Fellowship 100262Z/12/Z, a grant from KTRR and Human Frontiers Science Program Research Grant RGP0033/2015 to M.L.D., and a Wellcome Trust Senior Research Fellowship (207537/Z/17/Z) for O.D.

## Materials and Methods

### Reagents

RPMI 1640 (31870074), was purchased from ThermoFisher Scientific. Hyclone Fetal bovine serum was obtained from Fisher Scientific. The following anti-human antibodies were purchased from Biolegend, CD62L (DREG-56), CD3 (OKT3), CD8 (HIT8a), HLA-A2 (BB7.2), CD80 (2D10), CD86 (IT2.2), CD40 (5C3), CD69 (FN50), 4-1BB (4B4-1), CD107 (H4A3), TNFα (Mab11), and anti-mouse TCRβ (Η57-97). The following anti-human antibodies were purchased from BD Bioscience, CD45RA (HI100), CD4 (RPA-T4), and CD11c (B-ly6), CD14 (MΦP9) and PD-1 (EH12.2H7). Nivolumab (Ab00791-13.12) was obtained from Absolute Antibody. The following anti-human antibodies were purchased from eBioscience CD83 (HB15e) IFNγ (4S.B3) and PDL-1 (MIH1). HLA-A*0201 with 9V was generated and folded in house. HLA-DPB1*04:01 biotinylated monomers were obtained from the NIH tetramer facility. NY-ESO-9V_157-165_ (SLLMWITQV), NY-ESO-4D_157-165_ (SLLDWITQV), NY-ESO-6T_157-165_ (SLLMWTTQV) and NY-ESO-9L_157-165_ (SLLMWITQL), MAGE-A3_243-258_ (KKLLTQHFVQENYLEY), NY-ESO_161-180_ (WITQCFLPVFLAQPPSGQRA), HIV p17 GAG_77-85_ (SLYNTVATL) were purchased from GeneScript with > 95% purity and resuspended in DMF at 1mg/ml and stored at −20°C.

### Plasmids and constructs

The sequences for the 1G4 TCR were provided by Vincenzo Cerundolo, the 868 TCR was provided by Andrew Sewell, and all the MHC-II restricted TCRs (SG6, R12C9 and 6F9) were shared by Steven Rosenberg.

All TCRs were synthesised using GeneArt services from ThermoFisher, by adding Kozak sequence and codon optimisation for human expression. The 1G4 and 868 were to include a cysteine modification in the alpha and beta chains (see Supplementary table for sequences).

The MHC-II restricted TCRs were synthesised so that the human constant region was replaced with the mouse constant regions (see Supplementary table for sequences).

The alpha and beta chains were cloned separately into a pGEM-GFP-64A plasmid (gift of James Riley) between HindIII and NotI for MHC-I restricted TCRs and for human CD3ζ and between AgeI and NotI for MHC-II restricted TCRs.

### T-cell purification

T-cell were isolated from anonymised leukopoiesis products obtained from the NHS at Oxford University Hospitals by (REC 11/H0711/7). Resting human T-cell subsets were isolated using negative selection kits (Stemcell Technologies), total CD4 or CD8 T-cells were enriched using RosetteSep kits from Stemcell Technologies, followed by EasySep kits for the corresponding naïve and memory sub-population, following the manufacturer’s protocol. Cell purity was assayed with anti-CD4, anti-CD8, anti-CD62L and anti-CD45RA and all cells used had >95% purity.

Human CD4 and CD8 T-cells were cultured at 37°C, 5% CO_2_, in RPMI 1640 (Roswell Park Memorial Institute) medium Supplemented with 10% FBS (Gibco), 5% Penicillin-Streptomycin, Penstrep (Gibco), 1x MEM Non-Essential Amino Acids Solution, 20 mM HEPES, 1 mM Sodium Pyruvate, 2 mM Glutamax and 50 μM 2-mercaptoethanol (Sigma) (all from Thermo Fisher unless stated otherwise).

For T-cell activation 1:1 anti-CD3/anti-CD28 human activation beads (11132D) were added at 1:1 ratio with 50U/ml of IL-2 for two days, followed by removal of the beads and propagation of the culture in IL-2 until day 7.

### Express monocyte-derived dendritic cell generation

Monocytes were enriched from the same leukopoiesis products as T-cells using RosetteSep kits (Stemcell Technologies), and were then cultured at 1-2×10^6^/ml in 12 well-plates with differentiation media containing 50 ng/ml Interleukin 4 (IL-4, 200-04 A, Peprotech) and 100 ng/ml granulocyte-monocyte colony stimulating factor (GM-CSF, 11343125, Immunotools) for 24 h. For maturation the following cytokines were added for an additional 24 h, 1 μM Prostaglandin E_2_ (PGE_2_, P6532, Sigma), 10 ng/ml Interleukin 1 beta (IL1β, 201-LB-025/CF, Biotechne), 50 ng/ml Tumour Necrosis Factor alpha (TNFα, 300-01A, Peprotech), and 20 ng/ml Interferon gamma (IFNγ, 285-IF-100/CF, Bio-techne). Cell purity of the monocyte population was assessed using antibodies against CD14 and CD11c and typically was above 80%. Any non-monocyte contaminants can be removed by adhering the cells for 2-3 hours immediately following isolation followed by washing any unbound cells and applying the differentiation media.

Peptide loading on DC was quantified using an AlexaFluor 647 conjugated high affinity soluble TCR (c113 against NY-ESO) ^26^ with a known dye ratio and Quantum MESF AlexaFluor 647 beads as calibration (647, Bangs Laboratories).

Secretion of soluble factors was assessed using 0.5 ml of culture medium, and the corresponding controls of media containing the differentiation and activation cytokines, following the manufacturer’s protocol (Proteom Profiler Human XL Cytokine Array kit, ARY022B, R&D Systems).

### HLA typing of donors

For antigen specific experiments the donors were assessed for the relevant HLA, for CD8 based experiments 50 μl of whole blood was taken for flow cytometry analysis using an anti HLA-A2 mAb (Clone BB7.2). For CD4 based experiments 50 μl of blood was used to extract DNA using QIAamp DNA Mini kit (51304, Qiagen). The product was used for PCR analysis using the AllSet+ Gold HLA DPB1 high resolution Kit (54070D, vhbio) to determine if expressing the suitable haplotype. We were also able to induce the expression of a single chain dimer of HLA-A*02 in moDCs from donors which were HLA-A*02 negative using the mRNA electroporation approach.

### Engineering antigen specific T-cells

To express TCR constructs within resting naïve and memory T-cells, we used mRNA electroporation. mRNA preparations of the relevant TCRα, TCRβ and CD3ζ chains from a linearised pGEM-64A vector or a T7 containing PCR product was done using mMESSAGE mMACHINE T7 ULTRA Transcription kit as per manufacturer’s protocol (ThermoFisher, AM1345). The mRNA was purified using MegaClear kit (ThermoFisher, AM1908) and was aliquoted and kept at −80°C ^27^. To achieve high efficiency of expression, mRNA quality was assessed using agarose gels and samples showing any sign of degradation were discarded. Any mRNA preparation with yields bellow 40 μg mRNA per 1 μg of DNA was considered of low quality, finally all mRNA aliquots were stored at concentrations > 1 μg/μl, which can be achieved by concentrating the mRNA product using ammonium acetate precipitation or by combining >3 reactions of *in vitro* transcription during the MegaClear clean-up step. T-cells were harvested and washed three times with Opti-MEM (LifeTechnologies). The cells were resuspended at 25×10^6^/ml and 2.5-5×10^6^ cells were mixed with the desired mRNA products and aliquoted into 100-200 μl per electroporation cuvette (Cuvette Plus, 2 mm gap, BTX). For each 10^6^ cells CD8 T-cells 5 μg of each TCRα, TCRβ and CD3ζ RNA was used. For 10^6^ CD4 T-cells 10 μg of TCRα and TCRβ was used. Cells were electroporated at 300 V, 2 ms in an ECM 830 Square Wave Electroporation System (BTX). The cells were then collected from the cuvette and cultured in 1 ml pre-warmed media. Amaxa and Neon electroporation systems were also tested but failed in either achieving efficient TCR expression or affected T-cell motility. For Amaxa electroporation, the human T-cell nucleofector kit (VAPA-1002) was used and the manufacturer protocol was followed. Three different settings were tested: V-24, U-14 and T-23. For Neon electroporation, 2.5×10^6^ cells were resuspended in 100 μl tip, and two settings were tested (1700 V 10 ms, 4 pulses and 2150 V, 20 ms 1 pulse) ^28^.

Note, extended culture (>5 days) prior to electroporation also results in marginal reduction in TCR expression efficiency, although not observed for control GFP.

The exogenous TCR can be detected up to 96 h post electroporation.

### Supported lipid bilayers (SLB)

SLBs were prepared as previously described ^29^. In brief, piranha and plasma cleaned coverslips were mounted on sticky-Slide VI0.4 chamber (ibidi). Small unilamellar liposomes (LUVs) were prepared using 4 mM 18:1 DGS-NTA(Ni), “NTA-lipids”, (790404C-AVL, Avanti Polar Lipids,), 4 mM CapBio 18:1 Biotinyl Cap PE (870282C-AVL, Avanti Polar Lipids), “CapBio-lipids”, and 4 mM 18:1 (∆9-Cis) PC, “DOPC-lipids”, (850375C-AVL, Avanti Polar Lipids,). NTA-lipids were used at a final concentration of 2 mM and CapBio lipids were used at a dilution resulting in site density of 100 molecules/μm^2^. SLBs were allowed to form by incubating the coverslips with the appropriate LUVs for 20 min at room temperature (RT) followed by a wash with HEPES buffered saline (HBS) Supplemented with 1 mM CaCl_2_ and 2 mM MgCl_2_, and human serum albumin (HBS/HSA). The SLBs were loaded with saturating amounts of AlexaFluor568 labelled streptavidin, 5 μg/ml, (S11226, ThermoFisher) for 10 min at RT followed by pMHC at 100 molecules/μm^2^ and ICAM-1 200 molecules/μm^2^ for 20 min at RT.

### Micro-contact printed chambers

Micro-patterned surfaces presenting pMHC molecules were prepared using micro-contact printing by modifying the procedures described previously ^13^. Topological masters with repetitive circle patterns of 10 μm in diameter, spaced 30 μm centre-to-centre on a square grid were used to cover the entire channel of sticky-Slide VI^0.4^ chamber (ibidi). The master was used for casting of polydimethylsiloxane (PDMS) elastomer stamps. Sylgard 184 (Dow Corning) was used by mixing 1/7 (v/v) of curing agent to the elastomer. Rectangular stamps of PDMS were coated with biotinylated, AlexaFluor 674 labelled, Fc IgG (AG714, Merck Millipore) at 2 μg/ml in 150 μl of PBS for one hour. The blocks were then rinsed extensively in PBS, PBS with 0.05% Tween-20 and finally in MilliQ-grade water followed by gentle drying with N_2_ to remove droplets of water. Fc coated PDMS blocks were stamped onto the cleaned coverslips for 5 minutes under ~20 g of load for efficient transfer. Attempts to directly stamp pMHC, avidin or streptavidin resulted in partially or completely non-functional proteins. Streptavidin stamps were also non-uniform and eroded with subsequent washes. The patterned coverslip was then affixed to the sticky-Slide VI^0.4^ and washed sequentially with MilliQ-grade water and PBS, then coated with 13.5 μg/ml of CCL21 in 30 μl for one hour, followed by 3 μg/ml of ICAM1 in 180 μl for 3 h at 37°C. Coverslips were then used immediately for the following steps or kept with PBS at 4°C overnight. The stamped and protein coated channels were blocked with 10% dialysed FCS (to remove any free biotin) for 30 min at RT. Then unlabelled streptavidin was introduced into the channel at 2 μg/ml for 10 min at RT, washed and followed by 9V/A2 pMHC at 1 μg/ml for 20 min at RT.

### Collagen gel chamber assembly

DC were loaded with the mentioned peptide for 90 min at 37°C. For imaging experiments T-cells were labelled with a volumetric dye as described below (*Imaging*) or loaded with cell trace violet (C34557, ThermoFisher) for proliferation assays otherwise kept unlabelled.

Collagen mix was made by modifying the approach of Gunzer et al ^18^ using 5 μl 7.5% sodium bicarbonate, 10 μl 10x MEM, 75 μl 3 mg/ml Bovine Collagen I (PureCol) and a chemokine, 1 μg/ml of CCL21 (300-35-100, Peprotech) or CCL19 (300-29B, Peprotech) or 0.3 μg/ml of SDF1-a/CXCL12 (300-28A, Peprotech). All the work with collagen was done on ice to prevent polymerisation. Other matrices tested: VitroCol Human Collagen I (5007-A, CellSystems), Matrigel (at final conc. of ~4 mg/ml, two batches tested with similar results), and GelTrex (A1413301, ThermoFisher, at final concentration of ~8 mg/ml, two batches tested with similar results). All 3D cultures were made with Bovine Collagen I, except for the comparison shown in Figure 4-Supplement 1B, and mainly with CXCL12 unless otherwise stated.

Cell containing media (15 μl) supplemented with 10% Human serum (S-101B-FR, APS) (or 10% FBS) is added to 35 μl of the collagen mix so that the final culture contained 200,000 CD8, 200,000 CD4s T-cells and DC at varying ratios (typically 1:1, 1:5 and 1:10 to T-cells). Lower cell numbers can be used for functional experiment but the reported numbers here are ideal for good density for imaging.

30μl of the collagen is then added to a μ-slide angiogenesis chamber or sticky-Slide VI^0.4^ chamber (ibidi) and put upside down in a 37°C incubator with 5% CO_2_ for ~60 min (to entrap the cells in the gel as it polymerises). The wells are then topped up with 30 μl media containing the same concentration of chemokines as the gel (for μ-slide angiogenesis) or 100 μl per well (for sticky-Slide VI^0.4^). For in gel loading experiments, the media contained a peptide at 2x the reported concentration. Samples are either taken for imaging or kept in the incubator for functional analysis. For PD-1 blockade, the antibody or isotype control was added to the T-cell for 15 min prior to incorporation in the gel so the final antibody concentration was 10 μg/ml.

Extended culture in the 3D system leads to ~50% reduction in cell speed due to chemokine desensitisation and can be overcome by the addition of fresh chemokine after 18 h.

### Flow cytometry

**1G4** TCR expression was assessed using 9V-HLA-A2 tetramers, **868** TCR expression was assessed using SL9-HLA-A2 streptamers (6-7004-001, iba) and **6F9**, **SG6** and **R12C9** expression was assessed with anti-mouse TCR β (H57) antibody. The MHC II restricted TCRs seem to have very low affinity towards their antigen preventing detection using tetramers (attempts to measure affinity using SPR suggest a Kd bellow 500 μM). In brief, 50,000 cells were stained with 2.5 μg/ml tetramer (9V/A2) or 5 μg/ml streptamers (SL9/A2) or 1 μg/ml antibody for 20 min at 4°C, washed with PBS containing 2% FBS and 2 mM EDTA and taken for analysis on a flow cytometry machine (FORTESSA X-20, BD).

To analyse samples from collagen gels, the gels were first digested into solution using collagenase VII (Sigma #C0773) at 100 U/ml for 1 h at 37°C. The supernatant subsequently removed, and the cells stained with the desired antibody combination for 20 min 4°C. For intracellular staining, the cells were cultured in collagen gels and Brefeldin A or monensin were added, after 20 h for 2 h to prevent cytokine secretion to the supernatant and retain them inside the cells, along with a stimulation of Phorbol 12-myristate 13-acetate (PMA, P8139, Sigma) at 1 μg/ml and ionomycin (I0634, Sigma) at 1 μM, the cells were then stained using the FoxP3 intracellular staining kit (00-5523-00, eBioscience) and the desired antibodies. For experiments measuring LAMP1 expression, an antibody against it was added at the start of the culture.

LDH Cytotoxicity Detection kit (MK401, Takara Bio) was used as per manufacturer instructions to detect cell killing.

### Imaging

Cells are labelled with either one of the volumetric dyes, CMFDA (250 nM, ThermoFisher), CMRA (500 nM, ThermoFisher) or Deep Red (100 nM, ThermoFisher) for 20 min in complete media. For imaging CD69 activation the collagen mixture described above was supplemented with a BV421nlabelled anti-CD69 at 1 μg/ml (adjustments in volume are made to the media to maintain the same final collagen concentration). For calcium imaging T-cells were loaded with 1 μM of Fluo4-AM (ThermoFisher) for 20 min at 37°C. Cells were imaged using either a Dragonfly Spinning Disk system, a Perkin Elmer Spinning disk or an Olympus FluoView FV1200 confocal microscope using a 30x Super Apochromat silicone oil immersion objective with 1.05 NA. All microscopes included an environmental chamber to maintain temperature at 37°C and CO_2_ at 5%. Time-lapse images were acquired every 30 or 60s. *z*-scans were acquired every 3 μm.

SLB imaging was performed on an Olympus IX83 inverted microscope equipped with a TIRF module and Photomertrics Evolve delta EMCCD camera using an Olympus UApON 150x TIRF N.A 1.45 objective.

### Data analysis

Microscopy data from collagen gels was rendered and analysed using IMARIS software (Bitplane) using built-in analysis tools. Speeds bellow 2μm/min were considered as stationary cells. Synapse images were analysed using Fiji. Image analysis for micro-contact printing data was done using TIAM ^30^ as described in ^13^. In brief, cells were tracked using DIC images. IRM and fluorescence signal-positive segments of the tracks were used to calculate the attachment rate and the number of cells interacting with stimulatory spots to extract the half-life of interactions where a simple first order rate was used dA/dt=−k[A]; where A is cell number and k is the off-rate constant representing the exit from the spots by dissolution of the IS. Data from IMARIS and FlowJo was plotted and analysed in Prism (GraphPad), where all statistical tests were performed.

## Supplementary Legend

**Figure 2- Supplement Figure 1.**
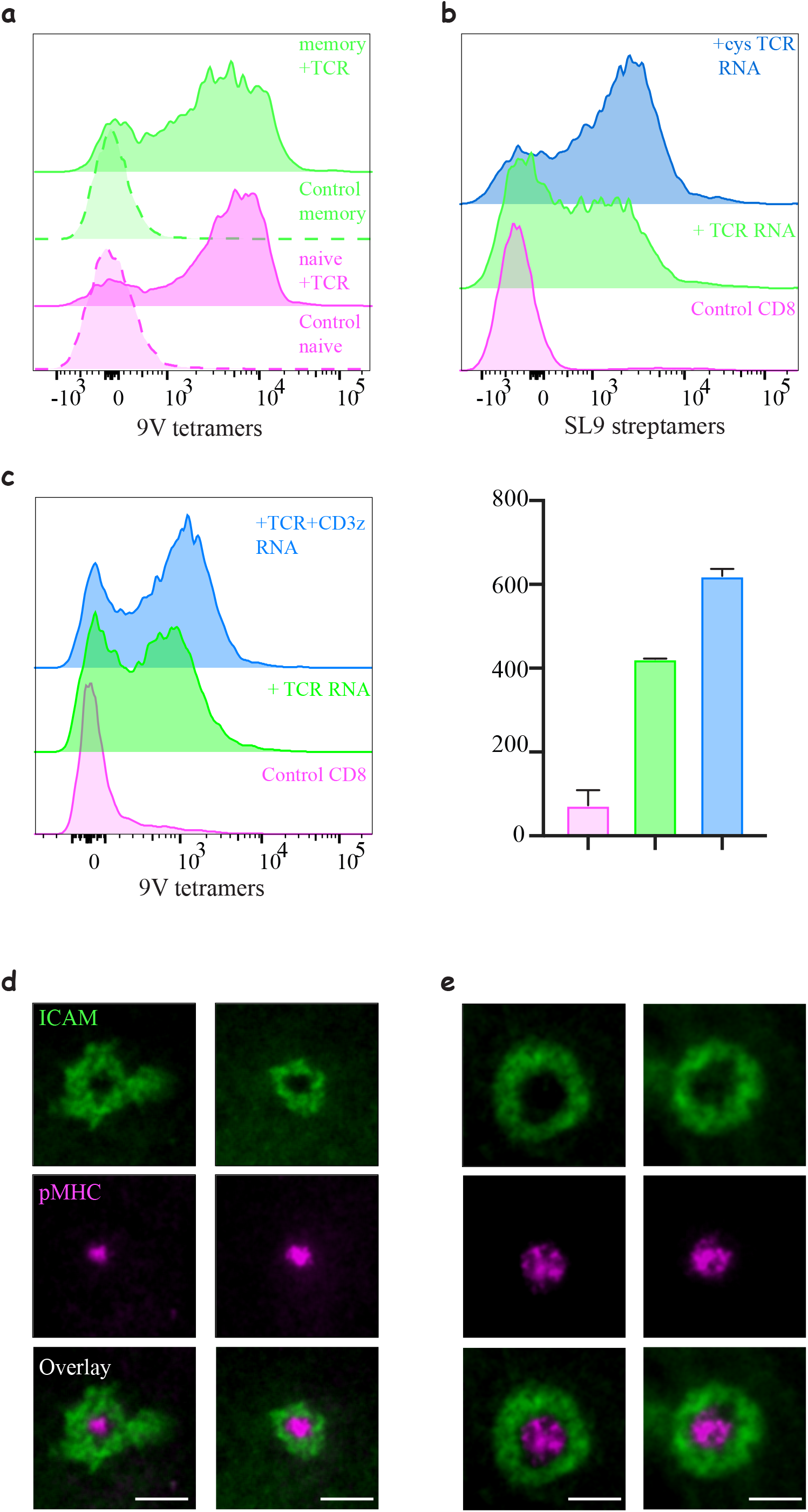
Engineering CD8 T-cells with different TCR constructs. (a) Expression of **1G4** in naïve and memory CD8 T-cells. Expression was comparable, although we noticed a slightly lower expression levels in memory T-cells. (b) Expression of **868** TCR using mRNA electroporation. Also note the improved efficiency after introducing the cysteine modification. Data shown are representative of at least 3 independent donors for the **868** without and with the cystine modification. Similar results were achieved for **1G4**. (c) Histograms of TCR expression following mRNA electroporating of the alpha and beta chain with or without the addition of CD3ζ(right) and quantification of the mean fluorescent intensity (left). Note the increase in efficiency (Representative of N>3). (d-e) Formation of an immunological synapse by **1G4** expressing naïve (d) and activated (e) CD8 T-cells on supported lipid bilayers (SLBs) with cSMAC enriched with 9V/A2 pMHC (magenta) surrounded by LFA/ICAM1 ring (green).

**Figure 2- Supplement Figure 2.**
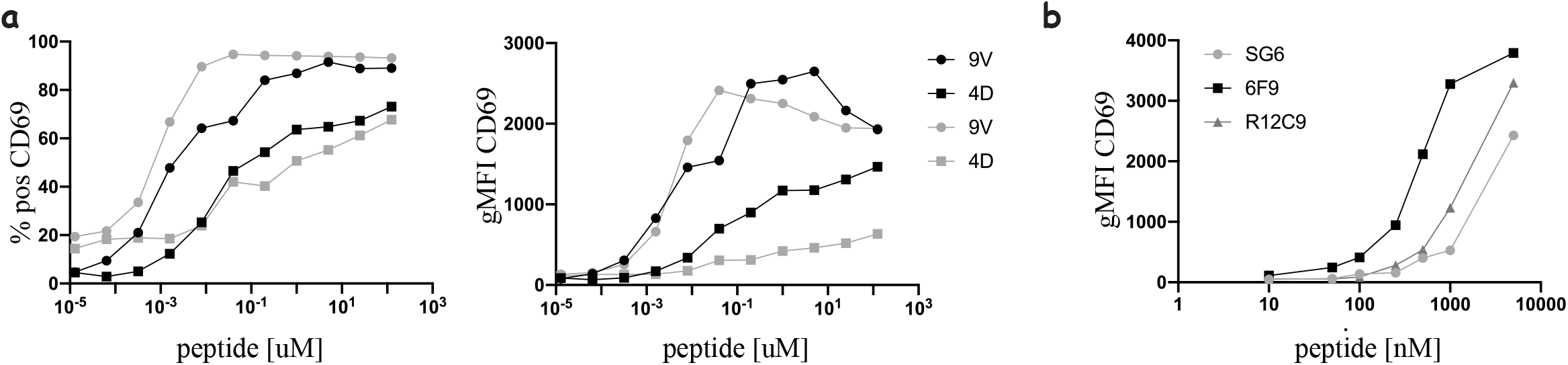
Functional response of mRNA electroporated T-cells. (a) Dose response curve of percent positive (left) and geometric mean (right) of **1G4**-expressing naïve (black) or memory (grey) CD8 T-cell upregulating CD69 upon stimulation with peptide loaded acDC, both with low (NY-ESO-4D) and high (NY-ESO-9V) affinity peptides. (b) Dose response curve of naïve CD4s expressing MHC-II restricted TCRs upon stimulation with peptide loaded acDC with either MAGE-A3_243-258_ (for 6F9 and R12C9) or NY-ESO_157-180_ (for SG6). T-cells and DCs are at 1:1 ratio.

**Figure 2- Supplement Figure 3.**
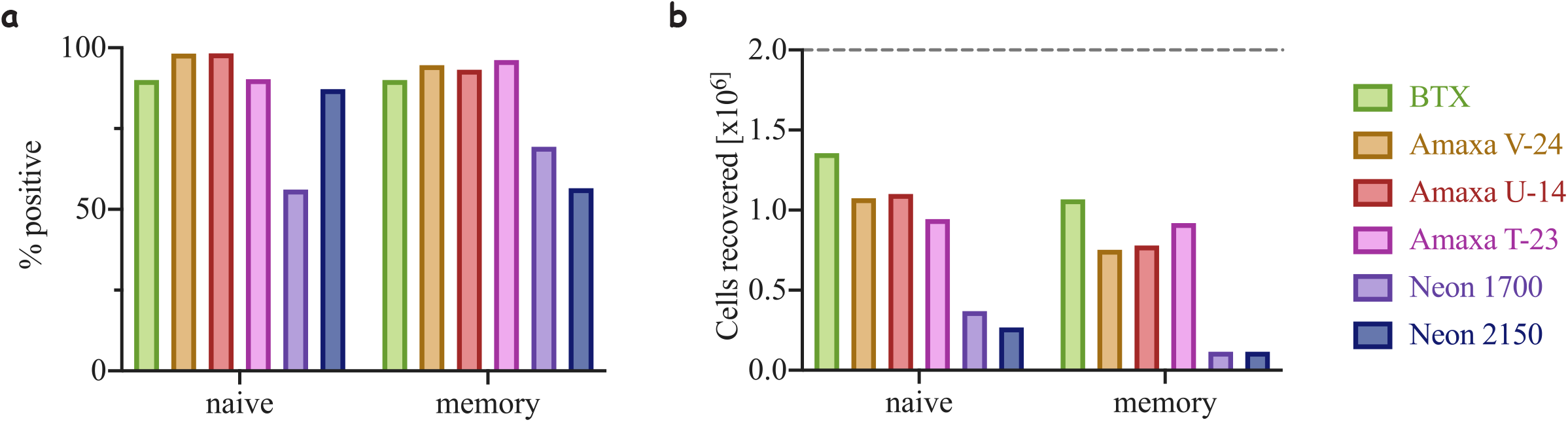
Efficiency of electroporation and cell recovery using different methods. (a) Percentage of Ruby positive cells following electroporation using three different commercially available electroporator: BTX, Amaxa from Lonza and Neon from ThermoFisher. U-14, V-23 and T-23 are electroporation setting on Amaxa. 1700 and 2150 are the voltages used on the Neon (see Methods for more details). (b) Number of cell recovered following electroporation- a product of cell viability and cell count compared to starting condition (dashed grey line).

**Figure 2- Supplement Figure 4.**
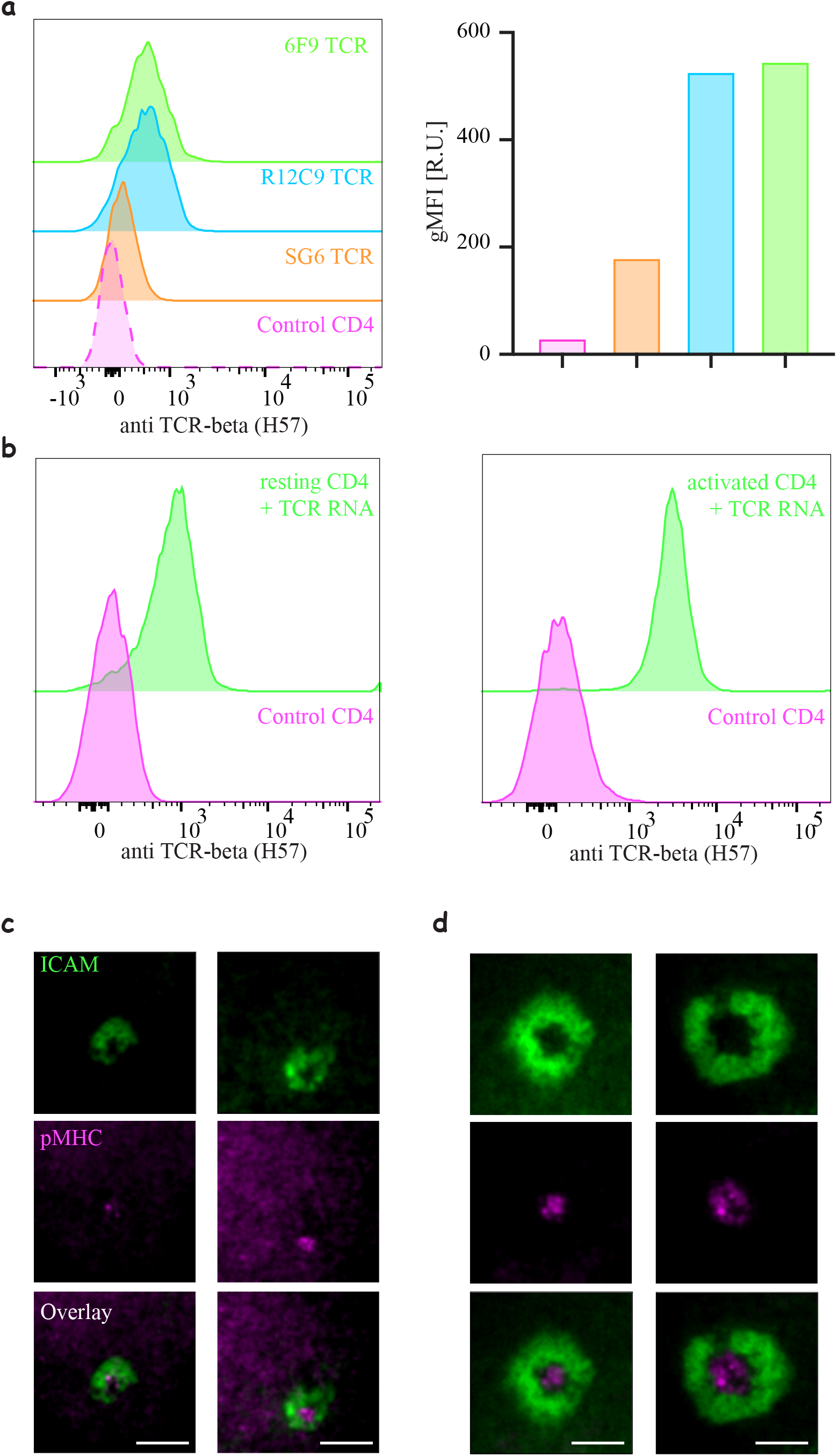
Engineering CD4 T-cells with different TCR constructs. (a) Histograms of TCR expression of three MHC-II restricted TCRs (**6F9**, **R12C9** and **SG6**) (right) and mean quantification of the mean fluorescent intensity (left). Note similar efficiency for **6F9** and **R12C9** compared to significantly lower efficiency of **SG6** (Representative of N>3). (b) Expression of **6F9** TCR by electroporation of naïve (left) or expanded (right) CD4 T-cells (similar results are obtained for **R12C9** and **SG6**). (c-d) Formation of an immunological synapse by **6F9** expressing naïve (d) and activated (d) CD8 T-cells on supported lipid bilayers (SLBs) with cSMAC enriched with 9V/A2 pMHC (magenta) surrounded by LFA/ICAM1 ring (green).

**Figure 2- Supplement Figure 5.**
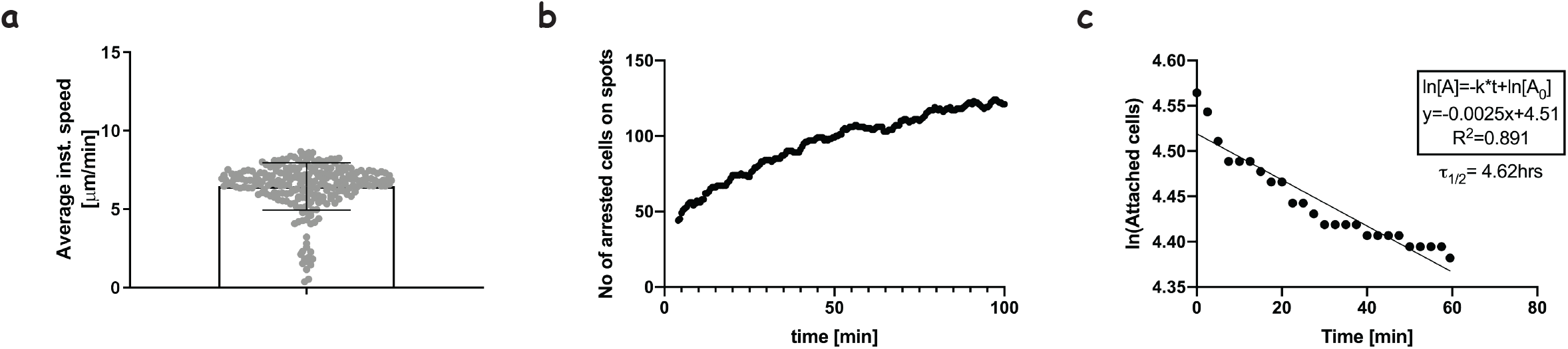
Interactions of 1G4 expressing naïve CD8 T-cells with pMHC presented on spatially segregated stimulatory spots. (a) The average speed of naïve CD8 T-cells moving on the micropatterned surfaces that have not yet attached to a stimulatory spot (Supplementary Video 1). This is essentially indistinguishable from the speed of untouched naïve CD8 T-cells in an analogous setting ^30^ (b) The number of cells arresting on spots as a function of time. Gradual accumulation of arrested cells is an indication of efficient search by the T cells and induction of arrest due to TCR-pMHC interactions. The attachment profile is comparable to that of untouched cells engaging with anti-CD3 stimulatory spots ^31^. (c) The number of remaining attached cells on stimulatory spots as a function of time after the first 40 minutes of imaging. The linear fit allows the extraction of the life-time of the interactions. Half-life of 4.6 hours compares well with that of untouched cells engaging with anti-CD3 spots ^13^.

**Figure 3- Supplement Figure 1.**
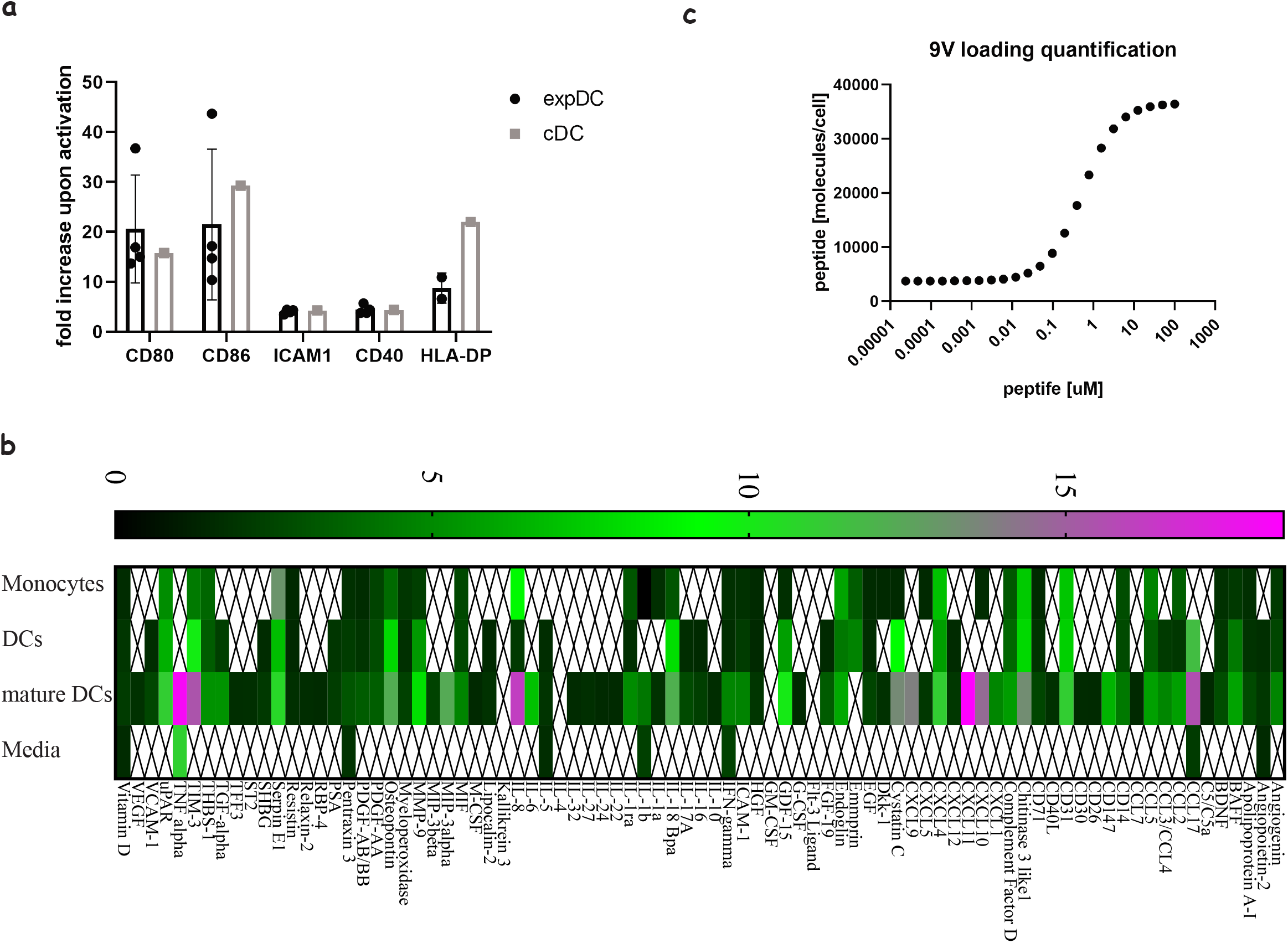
Comparing “express” and “classical” monocyte-derived dendritic cells. (a) The degree of costimulatory molecule upregulation upon maturation for classical and express DCs. (b) Analysis of fold production of cytokines and chemokines released from classical DCs generated in 7-day differentiation and maturation protocols at different stages: monocytes, differentiated DCs and matured/activated DCs. An average of three donors where signals bellow 1.5-fold above background were not included. (c) Quantification of the number of NY-ESO-9V peptides loaded on DCs using soluble high affinity TCR (**c113**) and MSEF calibration beads.

**Figure 4- Supplement Figure 1.**
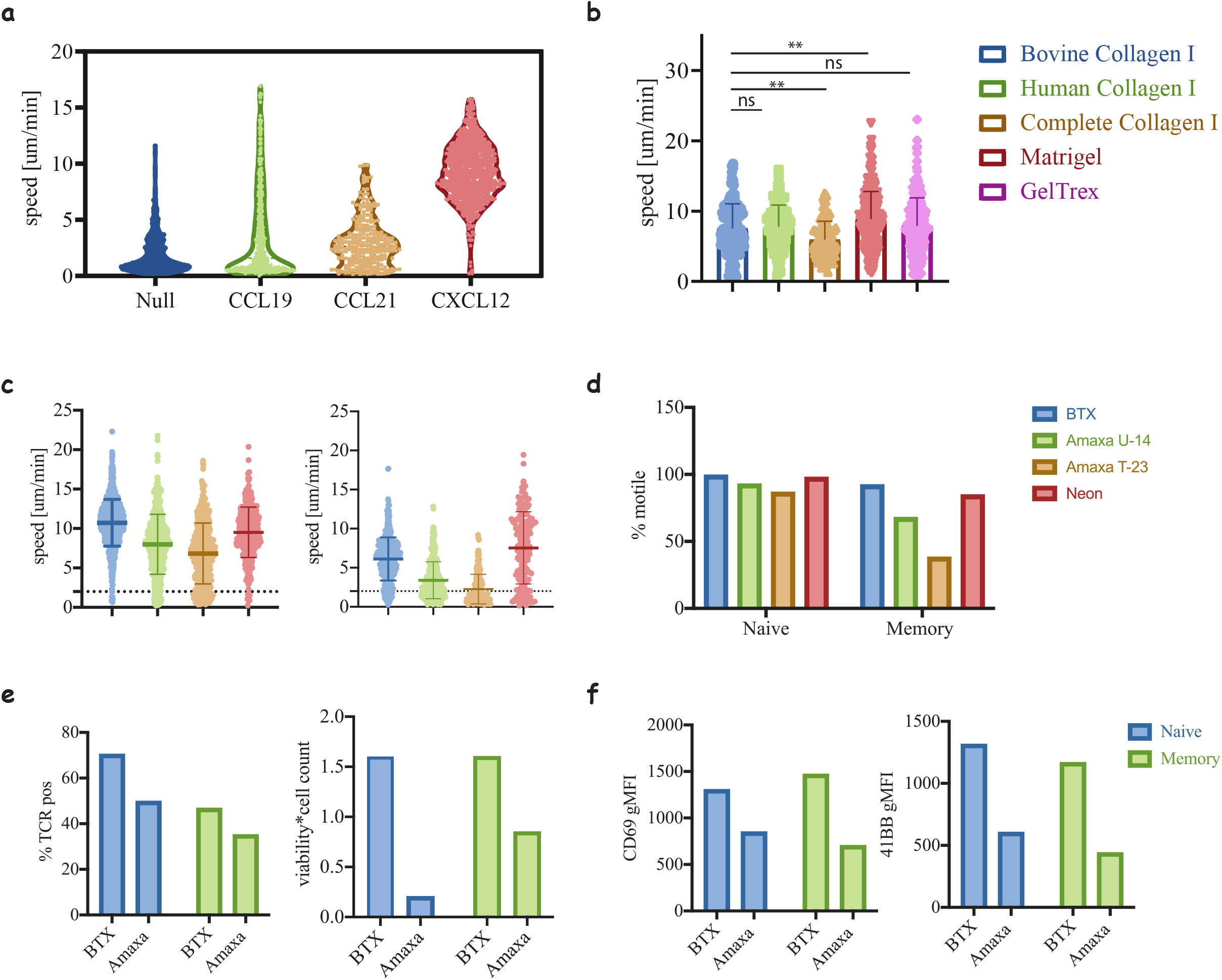
Motility of T-cells in 3D matrices. (a) Motility of naïve CD8 T-cells cultured in media containing FBS in the presence of different chemokines. Note that in the absence of human serum only the addition of CXCL12 supports cell motility (compared to Fig. 2a). (b) Motility of naïve CD8 T-cells in different matrices, collagen of different origins: human, bovine and non-tryptic bovine, and complex ECM: Matrigel and GelTrex (**, p < 0.01, ANOVA). (c) Motility speed of naïve (left) and memory (right) cells electroporated with different electroporation protocols (****, p < 0.0001, ANOVA, compared to BTX condition). (d) The percentage of motile cells was extracted from (c) to clarify the sensitivity of the cells to the electroporation method. (e) Comparing Amax and BTX electroporation for TCR expression (left) and cell recovery (left). (f) Measuring the activation of the cells from (e) after co-culture with peptide loaded acDCs (100nM NY-ESO-9V) in collagen gels at a ratio of 5:1 T-cells:DC. Data shown are representative of three repeats.

**Figure 4- Supplement Figure 2.**
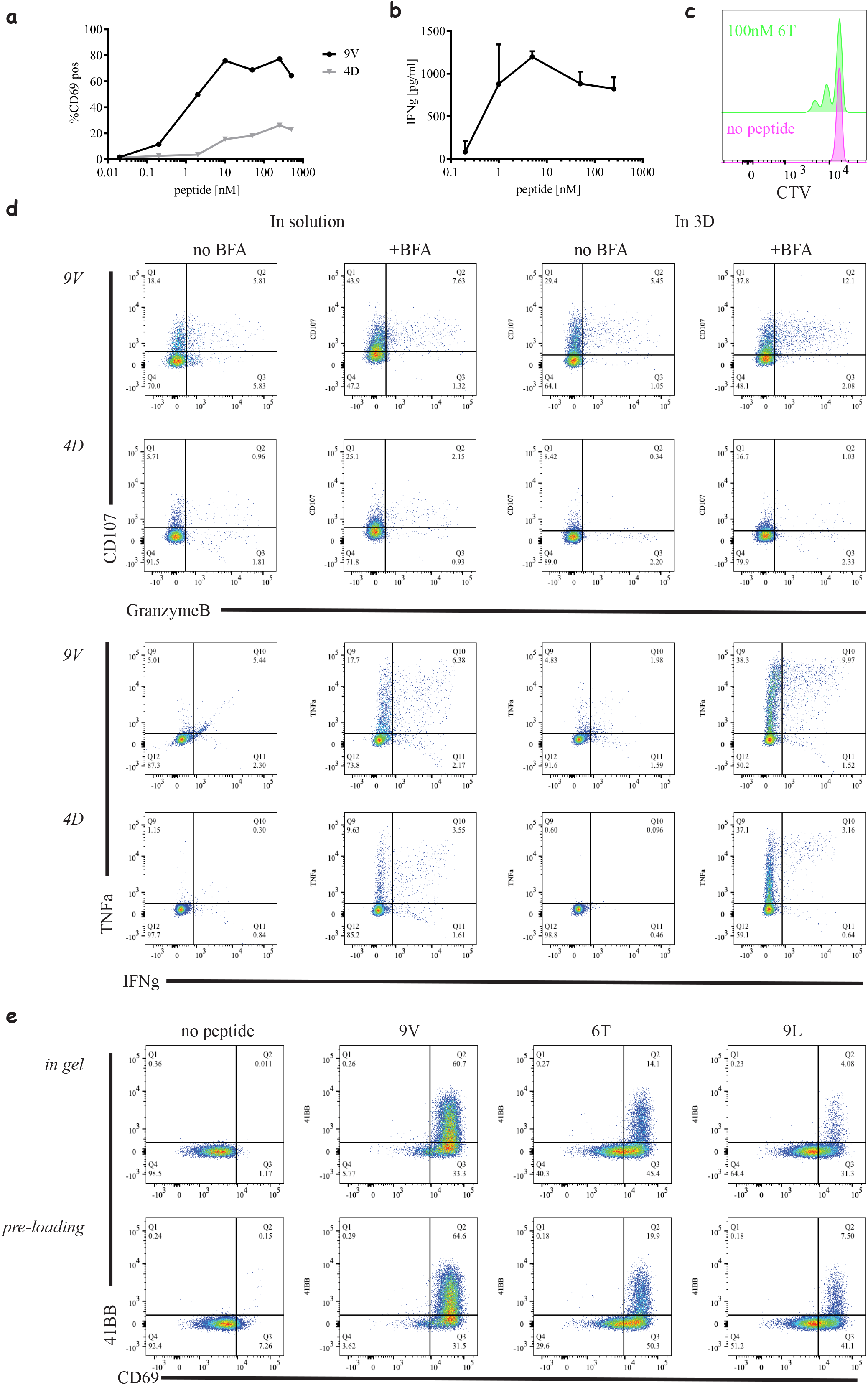
Different readouts in 3D. (a) Percentage of activated **1G4** - expressing CD8 T-cells as a function of peptide dose similar to Supplementary Fig. 2b, in collagen gels with 5:1 T-cells to DC. (b) Production of interferon gamma from a 24 h co-cultures of **1G4**-expressing CD8 T-cells and peptide loaded acDC with the indicated concentrations of NY-ESO-9V in collagen gels. (c) Proliferation of **1G4**-expressing naïve CD8 T-cells cultured with 100 nM NY-ESO-6T loaded acDC in collagen gels after 72 h, measured by cell trace violet (CTV) dilution. (d) Intracellular staining of activated **1G4**-expressing naïve CD8 T-cells stimulated with high affinity (50 nM, NY-ESO-9V) and low affinity (200 nM, NY-ESO-4D) loaded acDCs at 5:1 ratio, in solution and in collagen gels for comparison. The following markers were tested in the presence and absence of brefeldin A (BFA): Granzyme B, LAMP-1 (CD107), IFNγ and TNFα. (e) Activation of **1G4**-expressing naïve CD8 T-cells cultured in collagen gels with either pre-loaded acDCs or acDCs loaded inside the gel (Supplementary Videos 10 & 11 respectively).

**Figure 5- Supplement Figure 1.**
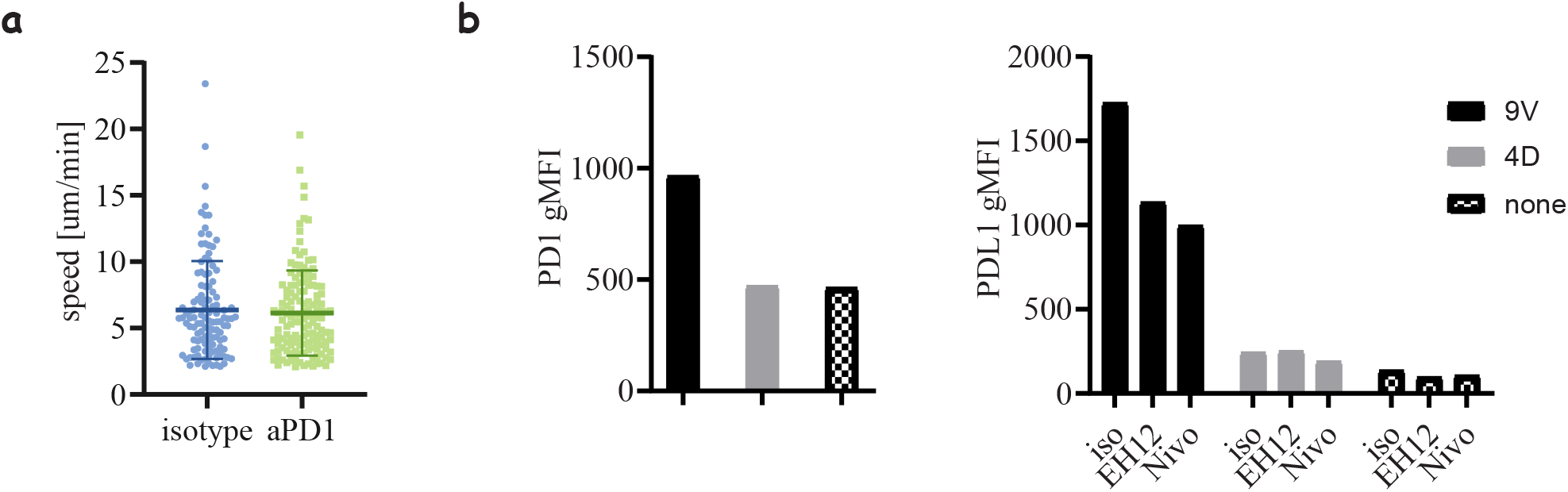
PD1 blockade effects on cell motility and regulation. **(a)** Motility speed of 1G4-expressing memory CD8 T-cells in co-culture with peptide loaded acDCs (100nM NY-ESO-9V) in the presence and absence of Nivolumab. (b) Expression of PD1 (left) and PDL1 (right) after stimulation with peptide loaded acDCs using high (100nM NY-ESO-9V) and low (100nM NY-ESO-4D) affinity peptides.

**Video 1**

A time-lapse movie showing gradual increase in the number of arrested cells on stimulatory spots presenting pMHC after some transient interaction events. The image series is a composite overlay of DIC, Interference reflection (IRM) and fluorescence images of the cells and microcontact printed FcIg (green) where cells form an IRM signal indicating spreading and durable interaction with the pMHC presented on stimulatory spots. Scale bar = each circular spot is 10 μm in diameter.

**Video 2**

3D reconstruction of naïve CD8 (magenta), naïve CD4 (cyan) T-cells, and Dendritic cells (green) moving in 3D collagen gels with RPMI containing human serum and exogenous human CXCL12.

**Video 3**

3D reconstruction of **1G4**-expressing naïve CD8 T-cells (magenta), interacting with antigen loaded acDCs (100 nM NY-ESO-9V, green) in 3D collagen in the presence of CCL19. Note the transition of cells between different DCs.

**Video 4**

3D reconstruction of **1G4**-expressing naïve CD8 T-cells (magenta), interacting with antigen loaded acDCs (100 nM NY-ESO-9V, green) in 3D collagen in the presence of CXCL12. Note the intermittent contacts and disengagement of T-cells before re-engaging with a different DCs.

**Video 5**

3D reconstruction of **1G4**-expressing naïve CD8 T-cells (magenta) interacting with 100 nM NY-ESO-9V loaded acDC (green) in the presence of irrelevant CD4 T-cells (cyan) and a soluble anti-CD69 (cyan). Following conjugate formation, upregulation of CD69 is observed by an accumulation of the antibody (cyan ring) around the CD8 T-cells (magenta).

**Video 6**

Maximum projection of 3D time-lapse of **1G4**-expressing naïve CD8 T-cells (magenta), loaded with calcium dye Fluo4-AM (green), interacting with antigen loaded acDCs (100 nM NY-ESO-9V, cyan) in 3D collagen and fluxing calcium (green) upon TCR/pMHC engagement.

**Video 7**

Maximum projection of 3D time-lapse of **1G4**-expressing naïve CD8 T-cells (magenta), interacting with acDCs (green) not loaded with NY-ESO antigenic peptide.

**Video 8**

Maximum projection of 3D time-lapse of **1G4**-expressing naïve CD8 T-cells (magenta), interacting with antigen loaded acDC (100 nM NY-ESO-9V, green).

**Video 9**

Maximum projection of 3D time-lapse of **1G4**-expressing naïve CD8 T-cells (magenta), interacting with antigen loaded acDC (100 nM NY-ESO-9L, green).

**Video 10**

3D reconstruction of **1G4**-expressing naïve CD8 T-cells (magenta), interacting with antigen loaded acDC (100 nM NY-ESO-6T, green).

**Video 11**

3D reconstruction of **1G4**-expressing naïve CD8 T-cells (magenta), interacting initially unloaded acDCs (green), where the peptide was later added after collagen polymerisation to a final concentration of 100 nM NY-ESO-6T.

**Video 12**

3D reconstruction of **1G4**-expressing naïve CD8 T-cells (magenta), **6F9**-expressing naïve CD4 T-cells (cyan) interacting with antigen loaded DCs (100 nM NY-ESO-6T and 10 μM MAGE-A3_243-258_).

